# Flow-Mediated Olfactory Communication in Honey Bee Swarms

**DOI:** 10.1101/2020.05.23.112540

**Authors:** Dieu My T. Nguyen, Michael L. Iuzzolino, Aaron Mankel, Katarzyna Bozek, Greg J. Stephens, Orit Peleg

## Abstract

Honey bee swarms are a landmark example of collective behavior. To become a coherent swarm, bees locate their queen by tracking her pheromones, but how can distant individuals exploit these chemical signals which decay rapidly in space and time? Here, we combine a novel behavioral assay with the machine vision detection of organism location and scenting behavior to track the search and aggregation dynamics of the honey bee *Apis mellifera* L. We find that bees collectively create a communication network to propagate pheromone signals, by arranging in a specific spatial distribution where there is a characteristic distance between individuals and a characteristic direction in which individuals broadcast the signals. To better understand such a flow–mediated directional communication strategy, we connect our experimental results to an agent–based model where virtual bees with simple, local behavioral rules, exist in a flow environment. Our model shows that increased directional bias leads to a more efficient aggregation process that avoids local equilibrium configurations of isotropic communication, such as small bee clusters that persist throughout the simulation. Our results highlight a novel example of extended classical stigmergy: rather than depositing static information in the environment, individual bees locally sense and globally manipulate the physical fields of chemical concentration and airflow.

## 1 Introduction

Animals routinely navigate unpredictable and unknown environments in order to survive and reproduce. One of the prevalent communication strategies in nature is conducted via volatile signal communication, e.g., pheromones [Conte and Hefetz, 2008, Kant et al., 2015]. As the range and noise tolerance of information exchange is limited by the spatiotemporal decay of these signals [Celani et al., 2014, Wilson et al., 2015], animals find creative solutions to overcome this problem by leveraging the diffusivity, decay, and interference with information from other individuals [Cardé and Willis, 2008, Bénichou et al., 2011, Gire et al., 2016, Hein et al., 2016].

For olfactory communication, honey bees use their antennae to recognize and respond to specific odors. Recent studies have revealed the bees’ distinct electrophysiological responses to different chemicals with quantifiable preferences [Zhao et al., 2020]. Olfactory communication with pheromones is crucial for many coordinated processes inside a honey bee colony, such as caste–recognition, regulating foraging activities, and alarm–broadcasting [Trhlin et al., 2011, Pankiw et al., 1998, Lensky and Cassier, 1995]. To become a coherent swarm in the first place, honey bees must locate their queen by tracking her pheromones that decay rapidly in time and space. How can honey bees that are far away from the queen locate her? The specific mechanisms of the signal propagation strategy are still unknown.

We propose that the mechanism to locating the queen involves a behavior called “scenting,” where bees raise their abdomens to expose the Nasonov pheromone gland and release the chemicals [Pickett et al., 1981, McIndoo, 1914, Peters et al., 2017] (Fig. 1A, Supplementary Movie S1 left panel). In the traditional chemical signaling chemotaxis and quorum sensing scheme the produced chemical signal is isotropic, as seen in early embryonic development [Cates and Tailleur, 2015, Bäuerle et al., 2018] and aggregation of amoebae in *Dictyostelium* [Brenner et al., 1998, Levine, 1994, Kessler and Levine, 1993]. On the contrary, scenting bees create a directional bias by fanning their wings, which draws air along the pheromone gland along their anteroposterior axis (Fig. 1A). We show that when bees perform the scenting behavior, they fan their wings to direct the signal away from the queen and towards the rest of the uninformed swarm. (Fig. 1B). This directional bias increases the probability that distant bees may sense those amplified pheromones, upon which they also stop at a certain distance from the scenting bee and amplify the signal along their own anteroposterior axis. The combination of detection and “rebroadcast” leads to a dynamic network that recruits new broadcasting bees over time, as the pheromones now travel a distance that is orders of magnitude the length of an individual. What are the advantages of a directional communication strategy vs. an isotropic one? And how do the bees harness the physics of directed signals to create an efficient communication network? To address these questions, we combined a novel behavioral assay, machine learning solutions for organism tracking and scenting detection and computational agent-based modeling of the communication strategy in honey bees and characterize the advantage of collective directional scenting.

**Figure 1:**
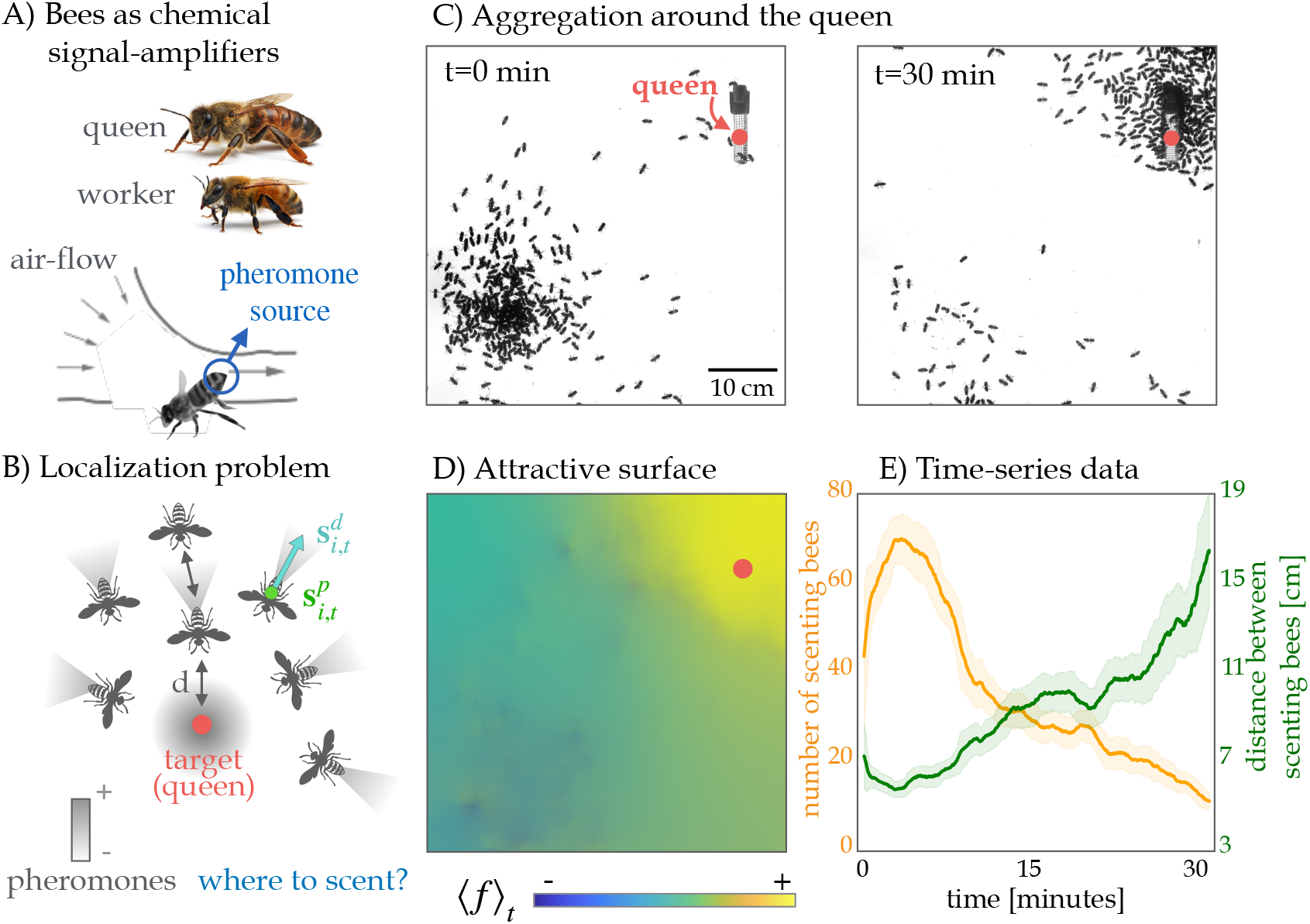
Honey bees use directed volatile communication to solve a localization problem of finding their queen. A) Pheromone sensing bees amplify the signal by opening the Nasonov gland in their abdomen, and fan their wings to transmit their pheromones backwards (a behavior called “scenting”). Photos of queen and worker bee were taken by Alex Wild; Scenting bee reproduced from [Peters et al., 2017]. B) Solution to the queen localization problem: by amplifying the queen’s pheromones, worker bees transmit the volatile signal over long distances while keeping a certain distance *d* from one another. 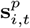 is the position of a given bee, and 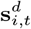 is the unit vector representing the direction of scenting. C) Example of a typical experiment where at *t* = 0 the queen is confined to a cage located at a corner of the arena, and the worker bees are located at the furthest corner. After ~ 30 minutes, the bees aggregate around the queen’s cage. D) The mean reconstruction of the attractive surface *f* according to Eq. S2, averaged over 30 minutes. The mutual information (Eq. S4) between this surface and the density of the bees at the end of the experiment illustrates the correlation between the scenting directions and the aggregation location of the bees. E) The average number of scenting bees and the average distance between scenting bees over time. The aggregation process is accompanied by a sharp increase and gradual fall in the number of scenting bees. In contrast, the distance between scenting bees is lowest at the peak in number of scenting bees at the beginning and gradually increases over time.

## 2 Behavioral Assay Results

To quantify the correlation between the scenting behavior of the bees, localization of the queen, and the aggregation process, we established a behavioral assay in which worker bees search for a stationary caged–queen in a semi–two–dimensional arena (see Supplementary Information section 5.1 for more details). We recorded the search and aggregation behavior of the bees from an aerial view for 1–2 hours (see Supplementary Movie S1 for a closeup example scenting bee with her abdomen pointed upwards and wing–fanning behavior in this experimental arena, in contrast with a non-scenting bee standing still). To extract data from the recordings, we then developed a markerless, automatic, and high–throughput analysis method using computer vision methods and convolutional neural networks (CNNs). This pipeline allows us to detect individual bees as well the positions and orientations of scenting bees (see Supplementary Information section 5.2 and Fig. S2 for more details, and Supplementary Movie S2 for the movie of the example in Fig. S2B,C).

### 2.1 Localization is correlated with scenting

In the 2–D experimental arena, worker bees placed at the opposite corner to the queen explore the space and eventually aggregate around the queen’s cage after ~ 30 minutes (Fig. 1C). For each scenting bee *i* at time *t*, we collected its position, 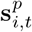, and direction of scenting, 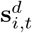 (unit vector). We then correlated the scenting events with the spatiotemporal density of bees in the arena by treating 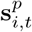 and 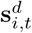 as a set of gradients that define a minimal surface of height *f* (*x, y, t*). This surface height *f* (*x, y, t*) corresponds to the probability that a randomly moving non–scenting bee will end up at position (*x, y*) by following the local scenting directions of scenting bees (see Supplementary Information section 5.4 for formal definitions and derivations).

To show that the attractive surface *f* (*x, y, t*) is correlated with the final concentration of bees *ρ*(*x, y, t*′), we compute the normalized mutual information, NMI(〈*f*〉_*t*_; 〈*ρ*(*t*_*end*_)〉), between the attractive surface averaged over the entire experiment (Fig. 1D) and the density of the bees averaged over the last 5 minutes of the experiment. The mutual information measures the information that the two variables, 〈*f*〉_*t*_ and 〈*ρ*(*t*_*end*_)〉, share, and determines how much knowing one variable reduces uncertainty about the other. The NMI is scaled between 0 (no mutual information) and 1 (perfect correlation). We averaged the density of bees over the last 5 minutes of the experiment to capture the density of the entire group and avoid discrete peaks of density resulting from movement of individual bees (see Supplementary Information section 5.4 for definitions). For the experiment shown in Fig. 1C–E, the NMI is 0.21. The top–right region in the arena with maximal 〈*f*〉_*t*_ corresponds to the location around the queen, illustrating the correlation between the scenting directions and the queen localization of the bees. We performed *N* =14 experiments of various group sizes, and report an average NMI of 0.11 (*σ*^2^ = 0.002). See Supplementary Information Table S1 for values for individual experiments and Fig. S3 for pairs of the average frame and the average attractive surface for all experiments.

### 2.2 A characteristic distance between scenting bees

The aggregation process is accompanied by a sharp increase in the average number of scenting bees, followed by a slow decay, until there are few scenting bees and the majority of the bees are aggregated around the queen (Fig. 1E, orange curve). See Supplementary Movie S3 for the experiment shown in Fig. 1C–E, with the attractive surface reconstruction and time–series data. We also characterize the temporal dynamics of the distance between neighboring scenting bees, which are treated as adjacent points in the Voronoi diagram for each frame (see example diagram in Fig. S2E–F and Supplementary Information section 5.3 for more details). Throughout the growth in the number of scenting bees, the distance between scenting bees decreases to a minimal value, then increases as the number of scenters decreases (Fig. 1E, green curve).

To show the reproducibility of the behavioral assay and assess the effect of group size in the aggregation process, we tested fourteen group sizes ranging from approximately 180 to 1000 bees. Three example experiments with *N*_bees_ = 320, 500, and 790 are shown in Fig. 2A and Supplementary Movies S4, S5, and S6. Each example includes three snapshots showing the process of aggregation around the queen’s cage over 60 minutes. The average number of scenting bees over time for all experiments of various group sizes are shown in Fig. 2B. Generally, there are more scenting bees over time as the number of bees in the arena increases. Across densities, we observe a typical characteristic of a sharp increase in scenting bees at the beginning as the bees initially search for the queen and a slow decrease as they slowly aggregate around the queen’s cage. Based on observations, we assume that bees in tight clusters, such as those at the very beginning, do not usually scent as they are not sufficiently spread out to fan their wings.

**Figure 2:**
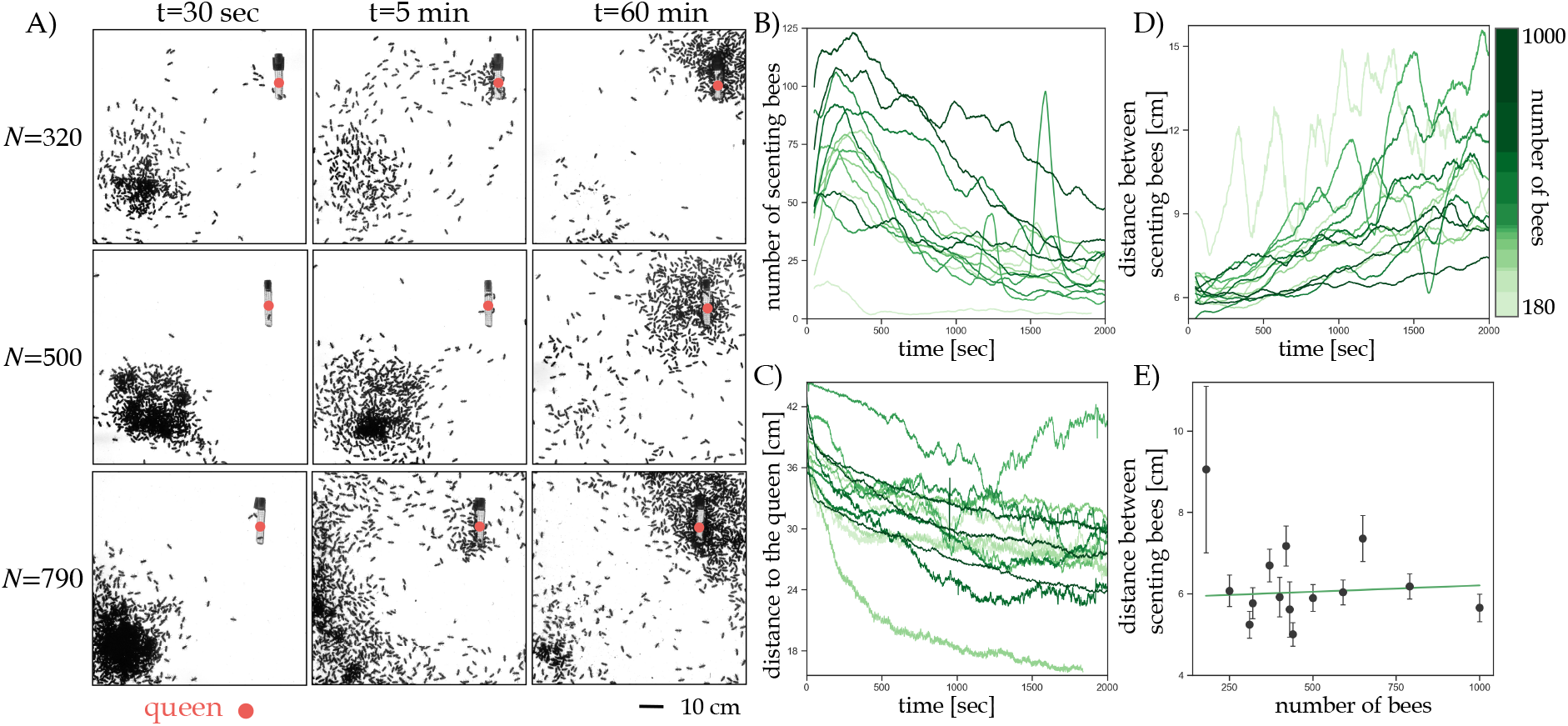
Distance between scenting bees is invariant to bee density. A) Example experiments of various bee densities, each with three snapshots showing the process of aggregation around the queen’s cage over 60 minutes. B) The average number of scenting bees over time for all experiments of various densities. There are more scenting bees with higher number of bees in the arena. Across densities, we observe a typical characteristic of a sharp increase in scenting bees at the beginning and decrease over time. C) The average distance to the queen over time for all experiments of various densities. This distance gradually decreases as bees aggregate around the queen. The positive correlation between distance to queen and number of scenting bees is evidence for the functional importance of scenting events to the problem of queen finding. D) The average distance between scenting bees over time for all experiments of various densities. Across densities, these distances are fairly constant at the beginning around the peaks in B), and increase over time. This is suggestive of a computational strategy that will be described in Section 3. E) The average distance between scenting bees at the peak in number of scenting bees as a function of density. Error bars represent the standard deviation over all the distances for each density. The distance between scenting bees is approximately constant across densities, except for a low–density outlier. A linear model (green line) is fit to the data, excluding the outlier, with a slope of *m* = 0.00031 cm/n_bees_.

The time evolution of the average distance from all bees in the arena to the queen is shown in Fig. 2C. There are no clear distinctions between the temporal dynamics of this distance as a function of density. For most densities, there is a sharp drop in this distance to the queen very early on (~ 100 --200 s), corresponding to the early increase in number of scenting bees. The average distance to the queen then gradually falls as bees make their way to her vicinity.

We also measured the distance between neighboring scenting bees over time for all group sizes (Fig. 2D). This distance shows a characteristic gradual increase over time. At the beginning of the aggregation process at the peaks in scenting bees, this distance is fairly constant across all sizes, ranging from 5.00 to 7.35 cm, with an outlier from the lowest group size experiment at 9.06 cm (Fig. 2E). A linear model is fit to the data (excluding the outlier) with the slope *m* = 0.00031 cm/n_bees_. The presence of this constant distance between nearest–neighbor scenting bees during the initial stage of the aggregation suggests the possibility of a pheromone concentration threshold that turns on scenting for individual bees in this collective communication network. As more bees aggregate around the queen, the bees collectively ‘turn off’ scenting. The higher distance between scenting neighbors later in the recordings suggest that the few remaining scenting events are more stochastic.

These experimental results, scenting behavior with wing–fanning to direct pheromones, the threshold-dependent triggering of this behavior, and a characteristic distance between scenting bees, serve as core ingredients for our agent–based model. We will use the model as a proof of concept of our proposal of the mechanistic localization behavior, described in more detail in the next section. The goal of our modeling is to test hypotheses of the mechanisms behind the phenomenon and explore the possible emergent patterns that arise to assess the effect of the directional signaling strategy employed by the bees.

## 3 Agent–Based Model Results

To identify the unifying behavioral principles that harness the dynamics of volatile signals, we developed a model that captures two important physical dynamics surrounding individual bees, substance diffusion and sensing local concentration gradients, while they perform search and identification. Our agent–based model is embedded in a flow environment where individuals can sense local concentrations of pheromones and propagate them backward as well as move uphill the gradient (see Supplementary Information section 5.5 and Fig. 3 for full model details).

**Figure 3:**
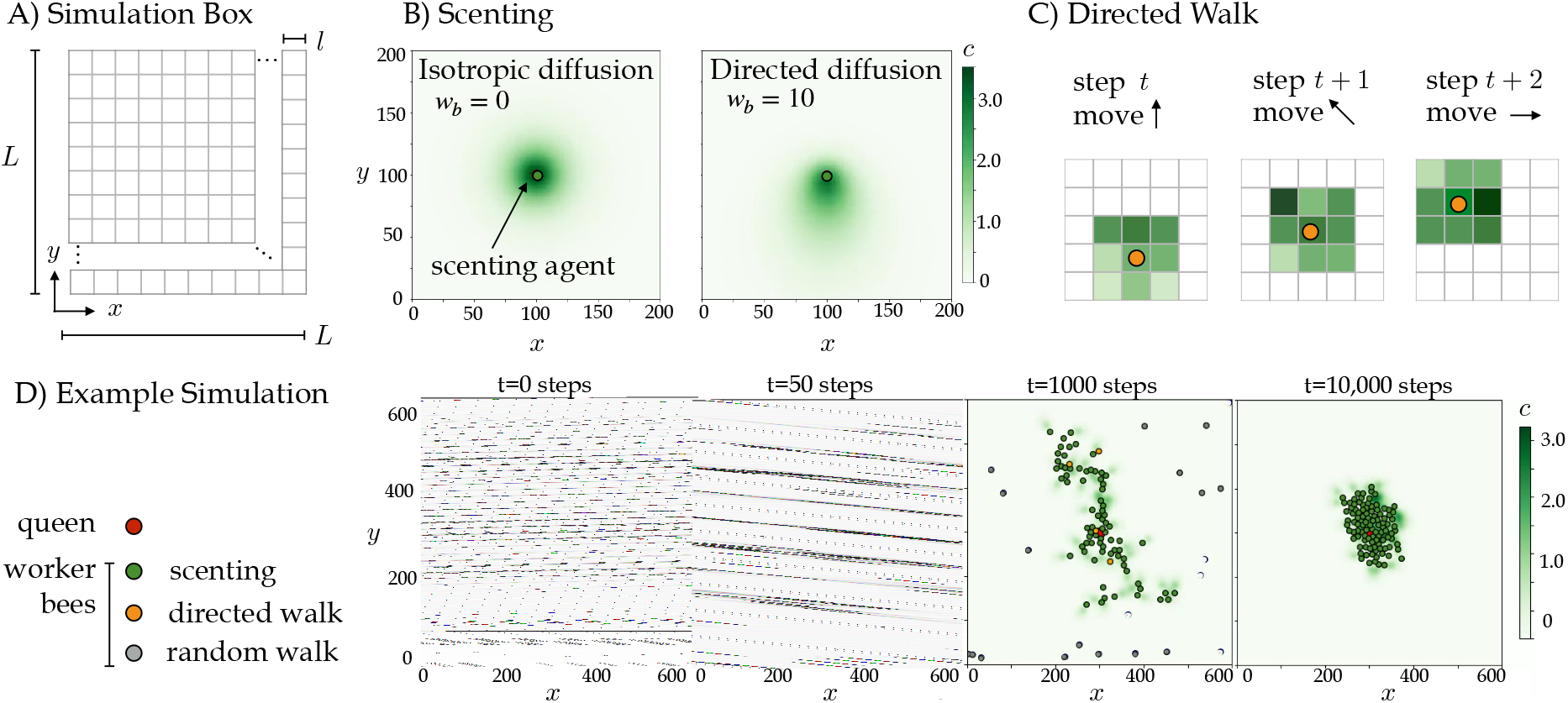
Scenting Model. A) Our *L* × *L* two dimensional simulation box is discretized into *l* × *l* sized pixels. B) Scenting bees can produce directional bias (example on the right with *w_b_* = 10), compared to no directional bias on the left. C) When a bee detects a local pheromone concentration that is above the activation threshold (*C*(*x, y, t*) > *T*), the bee calculates the concentration gradient around it (using the nearest 9 pixels, highlighted in different shades of green) and walks up gradient towards higher pheromone concentrations. D) Example of a simulation showing a series of snapshots. The queen is shown as a red filled circle, and worker bees are shown as a red filled circle colored by their internal state: scenting (green), activated (orange) and non–activated (gray). Activated bees perform a directed walk up the pheromone gradient, and non–activated bees are performing a random walk. The instantaneous pheromone concentration *C*(*x, y, t*) corresponds to the green color scale. Simulation parameters are *N* = 100, *w_b_* = 30, *T* = 0.01 *C*_0_ = 0.0575, *D* = 0.6, and γ = 108.

### 3.1 Model predicts optimal signal propagation within a range of behavioral parameters

We systematically explore a range of values for the directional bias *w_b_*, and the threshold *T*, for various numbers of bees in the arena *N*. Parameters *w_b_* and *T* are potential behavioral parameters that bees could adjust based on input from the environment. The directional bias *w_b_* represents the magnitude of the directional diffusion of pheromone released by a bee (see Fig. 3B for a comparison of isotropic (*w_b_* = 0) vs directed diffusion (*w_b_* = 10). The threshold *T* is the local concentration of pheromone above which a bee is activated from the random walk (i.e. a bee scents or walks up the concentration gradient when this threshold is met). Each combination of the these three parameters is a trial repeated 20 times. All other parameters, including *C*_0_ (the initial pheromone concentration), *D* (diffusion coefficient), and γ (decay constant) from Eq. S6, remain constant across all trials.

We quantify aggregation around the queen through the average distance of worker bees to the queen, 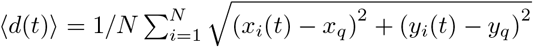, where *x_i_* (*t*) and *y_i_*(*t*) are the x and y positions of worker bee *i*, respectively, and *x_q_* and *y_q_* are the x and y positions of stationary queen bee. We also extract various other properties of the aggregation processes in the model: the queen’s cluster size and the number of clusters as additional measures of efficiency, the distance to the queen from the farthest active bee (i.e., a bee that is scenting or performing the directed walk up the pheromone gradient) to assess how far signals as a function of *w_b_* and *T* propagate, and the number of scenting bees as a measure of energy expenditure. Our model predicts 4 distinct phases for all densities, which are determined by (*w_b_, T*):

#### Phase 1

For low values of both *w_b_* and *T*, the bees aggregate into small clusters that are homogeneously spread throughout the simulation box (Fig. 4E). This is reflected by: 1) a sharp initial decrease and then gradual decrease of 〈*d*(*t*)〉 throughout the simulation (Fig. 4A), a consistently high number of scenting bees (Fig. 4B), a consistently small cluster of bees at the queen’s vicinity (Fig. 4C), and a consistently high distance of the farthest active bee from the queen (Fig. 4D). Simulation in Supplementary Movie S7.

**Figure 4:**
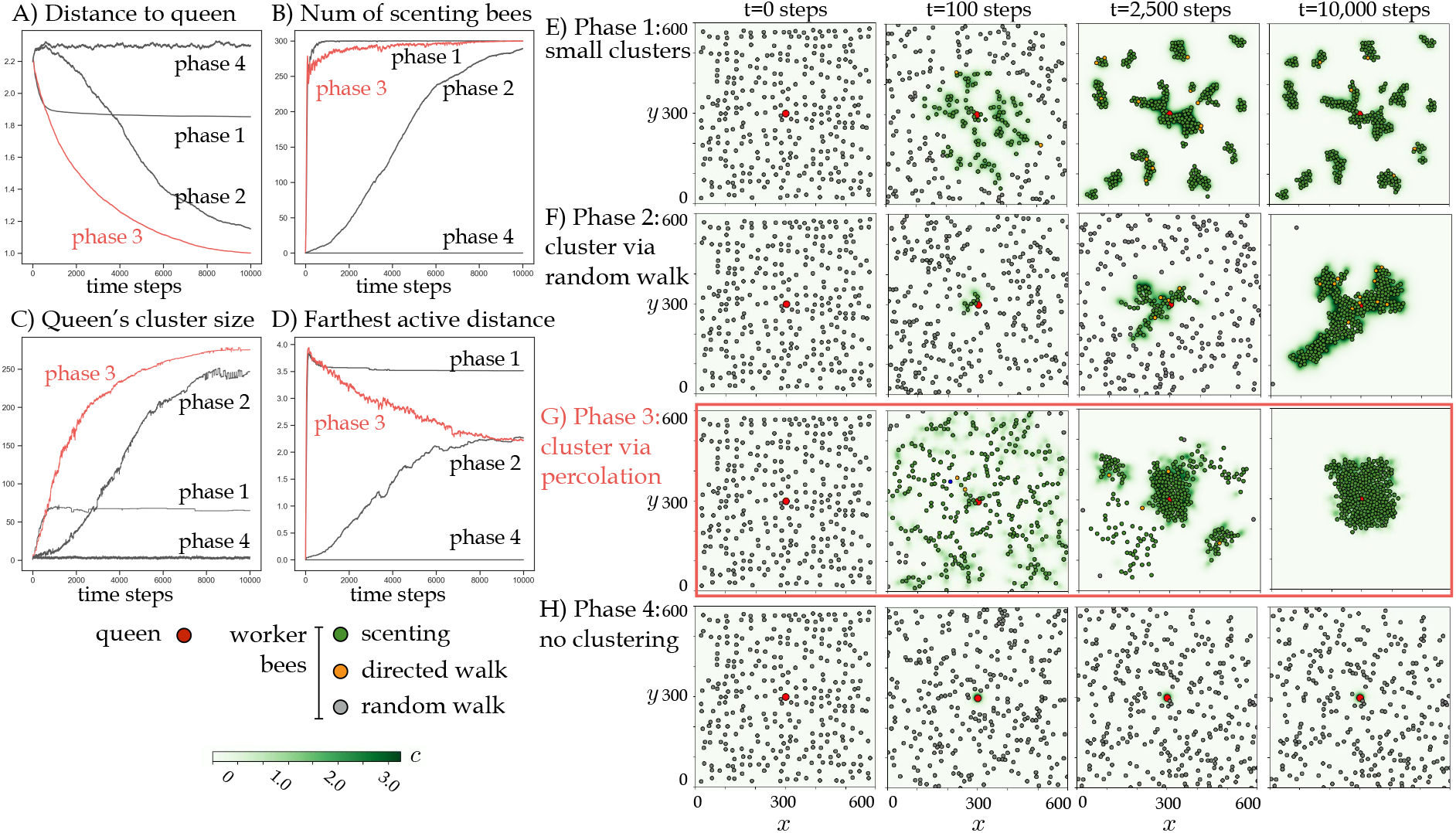
Directional bias is associated with optimal search and aggregation in the scenting model. A) The average distance of the worker bees to the queen as a function of time steps, for examples of the four different phases. B) The average number of scenting bees as a function of time steps, for examples of the four different phases. C) The average queen’s cluster size as a function of time steps, for examples of the four different phases. D) The average distance of the farthest active bee to the queen as a function of time steps, for examples of the four different phases. E–H) Example of a simulation from the four different phases, showing a temporal series of snapshots. The queen is shown as a red filled circle, and worker bees are shown as a red filled circle colored by their internal state: scenting (green), perform a directed walk up the pheromone gradient (orange) and performing a random walk (gray). The instantaneous pheromone concentration C(x,y,t) corresponds to the green color scale. Simulation parameters are *N* = 300, *C*_0_ = 0.0575, *D* = 0.6, and γ = 108. For phase 1 *w_b_* = 0, *T* = 0.01 (E); for phase 2 *w_b_* = 10, *T* = 0.5 (F); for phase 3 *w_b_* = 30, *T* = 0.05 (G); and for phase 4 *w_b_* = 0, *T* =1.0 (H). Phase 3, which is associated with the fastest aggregations around the queen, is highlighted in red.

#### Phase 2

At higher values of *T* and *w_b_*, only bees in the vicinity of the queen are activated and join the local cluster around the queen. This is reminiscent of Diffusion Limited Aggregation (DLA) that results in a sparse fractal–like cluster around the queen (Fig. 4F). This phase is reflected by a gradual decrease in 〈*d*(*t*)〉 and a gradual increase in the number of scenting bees, the queen’s cluster size, and the distance of the farthest active bee from the queen as bees slowly cluster via the random walk (Fig. 4A–D). Simulation in Supplementary Movie S8.

#### Phase 3

At low values of *T* and high values of *w_b_*, the activated bees create a percolating network of senders and receivers of the pheromone signal (Fig. 4G). This combination of *T* and *w_b_* ensures the fastest aggregation process around the queen and the fastest growth of the queen’s cluster (Fig. 4A, C). This process keeps most bees active with the scenting task throughout the simulation (Fig. 4B). Although bees in phase 2 and 3 eventually cluster at the queen’s location, the pheromone signals in phase 3 reach a much farther distance at the beginning than in phase 2 where bees slowly cluster only via the random walk (Fig. 4D). Simulation in Supplementary Movie S9.

#### Phase 4

When the activation threshold T is high enough, no worker bees are activated and no clusters are formed (Fig. 4H). In this phase, the bees simply perform a random walk. Simulation in Supplementary Movie S10.

The existence of phase 3 suggests that using a directional signal is advantageous, as this phase does not exist in the absence of directional bias (*w_b_* = 0). Although in both phase 2 and phase 3, the bees are able to aggregate around the queen with similar values of the average final distance to the queen (such as for *N* = 100, the values are 0.56 and 0.53 units, respectively, in Fig. S5D), bees in phase 3 are able to reach a plateau distance to the queen earlier (on average at *t* = 5117 compared to *t* = 6864.6 in phase 2, in Fig. S5E) while requiring less scenting events to reach that plateau (on average 5566.18 events compared to 7732.50 in phase 2, in Fig. S5F).

### 3.2 The effect of bee density on the phase boundaries

We use our model to construct phase diagrams for three different densities, low (*L*^2^/50), medium (*L*^2^/100), and high (*L*^2^/300). Across densities, low *T* and low *w_b_* result in phase 1 with multiple clusters (Fig. 5). In this phase, the range of *T* increases while the range of *w_b_* decreases as density increases. On the other hand, phase 4 is the result of high *T* and high *w_b_* values. With more bees in the arena, other phases expand while the range of phase 4 gets smaller.

**Figure 5:**
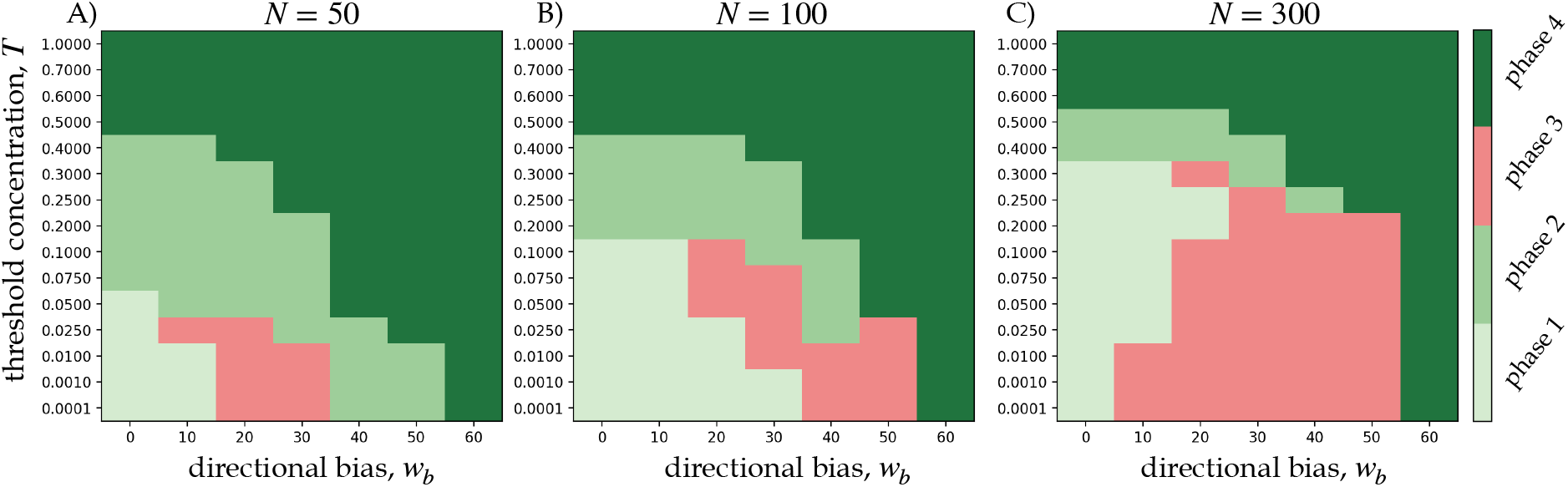
The effect of bee density on phase boundaries. Phase diagrams constructed from scenting model dynamics using summary heatmaps in S5 as a function of *T* and *w_b_* for all three densities, N=50 (A), N=100(B), and N=300(C). The area of the optimal phase (phase 3) grows with N.

Phase 2 is typically the result of intermediate *T* values and low to intermediate *w_b_* values. These higher *T* values prevent the percolation seen in phase 3, as the pheromone signals are not as far reaching when bees are less sensitive to sense them to become propagators. Phase 3 with the successful aggregation via percolation is achieved with intermediate to high *w_b_* and low *T* values. As density increases, the range of *w_b_* and *T* for phase 3 is greater as there are more bees to create and sustain the communication network in the arena. When there is higher density of bees, the bees are able to aggregate around the queen while not needing to be as highly sensitive (i.e., lower *T* value) to the signals. Across all densities, phase 3 never occurs when there is no directional bias (i.e., *w_b_* = 0). This illustrates the importance of the directed signaling strategy to create an effective communication network. The existence of a network of transmitters and receivers of pheromones that percolates across the entire simulation box is crucial to achieve fast and successful aggregation around the queen.

## 4 Discussion & Conclusion

Combining new experiments and high-throughput machine vision tracking of location and scenting behavior, we illuminated the communication mechanisms that honey bee swarms employ to locate their queen, a difficult problem given the limited information available from short–lived pheromone signals. We find that individual bees act as receivers and senders of signals by using the Nasonov scenting behavior, releasing pheromones from the glands and fanning their wings to direct the signals backward (Fig. 1A–B, Supplementary Movie S1 left pane). In an arena with a caged queen, bees were quick to activate a collective communication network, as we see a sharp, early–time increase in the number of scenting bees (Fig. 1E, 2B). In this network, scenting bees stand at a characteristic distance from their neighbors while dispersing signals (Fig. 2E), which suggests a concentration threshold in the activation mechanism of individual bees scenting behavior. We show that the scenting events are highly correlated with the collective aggregation around the queen (Fig. 1D, S3). Together, these experimental results provide testable hypotheses of the mechanisms of this collective communication strategy and of whether the threshold–dependent directional signaling behavior is advantageous – concepts that we explore with the agent–based model.

The experimental findings guide our design and implementation of an agent–based model of the queen–localization phenomenon. We model individual bees as virtual agents with relatively simple rules for movement, detection of pheromone based on a concentration threshold, directed signaling, and chemotaxis to move up the local gradient (Fig. 3). The virtual bees are not aware of the queen’s location or of the global pheromone profile in the flow environment. We show that by only local interactions, these virtual bees are able to aggregate around the queen mostly quickly and efficiently when they implement directed signaling within a range of bias values (i.e., 10 ≤ *w_b_* < 60) (Fig. 4). When the density of bees increases in the virtual arena, effective aggregation can also occur with a wider range of *T* as opposed to being limited to lower *T* or lower sensitivity to pheromones (Fig. 5). Thus with more bees active in the communication network, individuals can afford to be less sensitive while still achieving swarm formation. Overall, the four different phases that are present in the model show us the possible emergent properties of this honey bee phenomenon. Importantly, our modelling emphasizes the significance of wing–fanning behavior, which allows for directed signaling in the swarm.

In addition to presenting novel mechanistic details about the queen localization phenomenon in honey bees, we have also developed a new and effective image analysis pipeline for markerless, automatic, and high–throughput honey bee detection and behavior recognition (Fig. S2). Our approach uses state–of–the– art neural network models that are trained on our honey bee data and can be retrained and applied to other systems. Our high–throughput pipeline can be easily improved for future applications. First, the detection of individual bees currently employs a classical computer vision approach of simple Otsu’s adaptive thresholding, morphological transformations, and connected components. While these methods are quick and sufficient to identify bees from background, they struggle to separate bees when they significantly touch or overlap in the image, a problem exacerbated by our back light system. In future designs, we will modify our experimental setup to use different lighting systems where more features on bees are visible, allowing the usage of CNNs for the task of image segmentation to detect more individual bees in dense environment (e.g. [Bozek et al., 2018]). Additionally, while using the wing angle as a proxy for scenting in static images is effective, scenting is a time–dependent behavior which could be more accurately identified using information from multiple frames. Temporal information can be incorporated using activity recognition networks [Karpathy et al., 2014, Carreira and Zisserman, 2017] on labeled videos of bees. This will require tracking individual bees over time to build a labeled dataset composed of short movies of the scenting behavior. We will explore recent efforts in automatic tracking of bees, such as works by [Bozek et al., 2020]. The ability to track the scenting behavior of bees over time will allow us to answer interesting questions regarding the roles worker bees play in this swarming context. For example, does every bee eventually scent or does only a proportion of the same bees scent while others follow the pheromone trails to the queen?

Various future directions arise. The model will allow testable predictions about the resilience of the communication network. This concept includes assessing the effect of node failure via removal of some signaling bees, interference with a secondary signal via introducing artificial pheromone to the network, and the disruption of pheromone flows in the presence of wind. Experiments can then be performed to test the model’s phenomenological predictions. Together, the tools will allow better understanding of how the dynamic nature of the network allows the swarm to overcome local obstacles such as solid objects, deal with a non–stationary search target, as well as turbulent airflow and conflicting chemical signals.

From a physics perspective, our active system functions by coupling flows and forces in the presence of feedback [Peleg, 2019, Peters et al., 2019, Peleg et al., 2018]. The individual building blocks (in this case, a bee) can sense their micro–environment (flow, forces, chemical content) and respond in a way that promotes survival; typically, the response changes the macro–environment the individual is embedded in, thus creating a perpetual coupling between the individuals, the group, and the environment. From a biological perspective, our approach is an extension of the studies of classical stigmergy [Theraulaz and Bonabeau, 1999] wherein organisms deposit and respond to static information in the environment. In contrast, the bees in our system are able to sense local chemicals but also manipulate the global physical fields by actively directing signals with the scenting behavior. Harnessing the bees’ natural solutions to communication – honed by eons of evolution, selection, and refinement – we can not only more deeply understand collective animal behavior, but leverage that understanding to create bio–inspired system designs in the fields of dynamic construction materials, swarm robotics, and distributed communication.

## 5 Supplementary Information

### 5.1 Experimental setup

To quantify the correlation between the scenting behavior of the bees, localization of the queen, and the aggregation process, we established a behavioral assay and a high–throughput analysis method to automatically extract position and orientation information from the recordings.

European honeybees (*Apis mellifera* L.) are known to scent while standing or walking on solid surfaces, rather than flying. Hence, we restrict our indoors experimental arena to be semi–two–dimensional. The arena is composed of a wooden frame, and a plexiglass sheet, enclosing a volume of 50 cm x 50 cm x 1.5 cm. This space allows bees to walk and flap their wings, but keeps the bees from flying to avoid visual occlusions. To record the behavior of the bees, we mount a video camera (4k resolution, 30 fps) to a tripod, positioned directly above the arena to capture an aerial view (Fig. S1). We collect bees directly from our apiary at the BioFrontiers Institute. First, we capture a queen from our queen bank, and place her in mesh cage (10.5 x 2.2 x 2.2cm), long enough to provide moving space. This ensures that the queen is confined to the cage while worker bees can sense her pheromones and attend to her. We then collect roughly 2000 worker bees from one of our hives, and isolate them from their original queen for 24 hours. A dollop of fondant is placed in the bucket for the bees to feed themselves throughout their time inside the lab. After the queenless period, we introduce the workers to the caged queen and allow them to spend 24 hours together, for a total of 48 hours of adjustment before the experiments.

**Figure S1:**
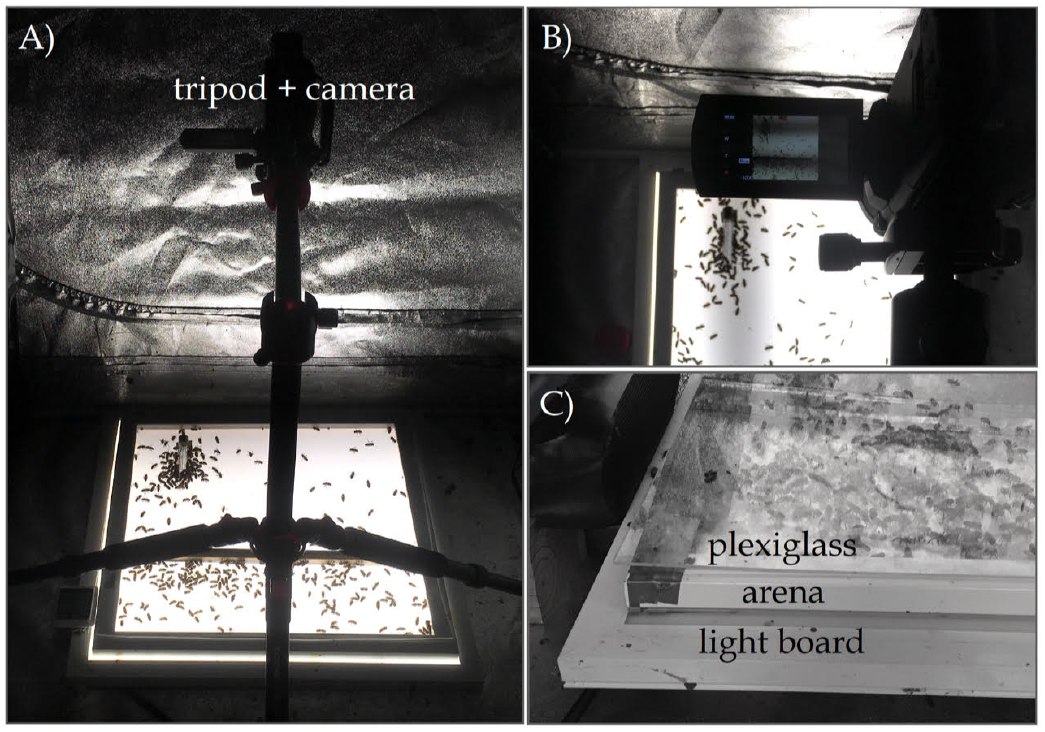
Setup for Behavioral Assay. A) Overview of setup, with a camera mounted on a tripod and positioned above the arena to capture an aerial view. B) Aerial view showing camera recording above the arena. C) Close–up of the arena, which is consisted of a light panel, the wooden frame as the arena on top of the panel, and a plexiglass sheet to enclose the experimental space.

Before the start of each experiment, the bees in the respective group are cooled down for approximately 5–10 minutes in a fridge (4° C) to deactivate the bees’ flying capability for easier handling. The queen bee in the mesh cage is isolated from the workers and placed into a position in the arena. Workers are then placed into the arena, and the plexiglass sheet is quickly placed on top to completely enclose the experimental space (Fig. S1C). Temperature of the arena is recorded from the beginning to the end of each experiment for a duration of 1 hour. Temperature is monitored regularly to ensure that the heat from the backlight board stays below 32 – −35°C and does not become a confounding factor to the fanning behavior. When the experiments are over, the bees are collected from the experimental area and returned to the apiary to their respective hive and queen bank. The arena and the plexiglass are cleaned (Lysol disinfectant wipes and wet paper towels) after each single experiment to remove any remaining pheromones for the next experiment.

### 5.2 Computer vision and deep learning approach for high–throughput analysis

To extract information from the experimental videos and characterize the collective aggregation via directed pheromone signaling, we employ various computer vision and deep learning approaches. These approaches allow us to perform automatic detection of individual bees and of scenting bees as well as estimation of the scenting direction.

We extracted color images from a given video at 30 fps and converted to 8-bit grayscale for storage, and used Otsu’s method to adaptively threshold the images, resulting in binary maps [Sezgin and Sankur, 2004]. Morphological opening is then applied to the binary images to erode then dilate them [Dougherty, 1992]. This process removes noise and separates individual bees from clustered groups. Connected components is applied to the binarized images, resulting in centroids (x, y positions) and pixel area. The pixel areas follow a multimodal distribution that represents junk artefacts (e.g., bee legs), individual bees, and larger bee groups that were not separated during the morphological transformations. The centroids and areas of individual bees are saved, and an additional processing step is performed on the large bee group detections. One more iteration of morphological opening is performed, followed by connected components. The resulting areas now follow a distribution where the smaller clusters represent individuals and show a sharp peak. The individual bee detections are added to the initial stack of individual detections, and the group detections are finalized as inseparable clusters. The centroid information is leveraged for cropping individual and grouped bees from the original images.

**Figure S2:**
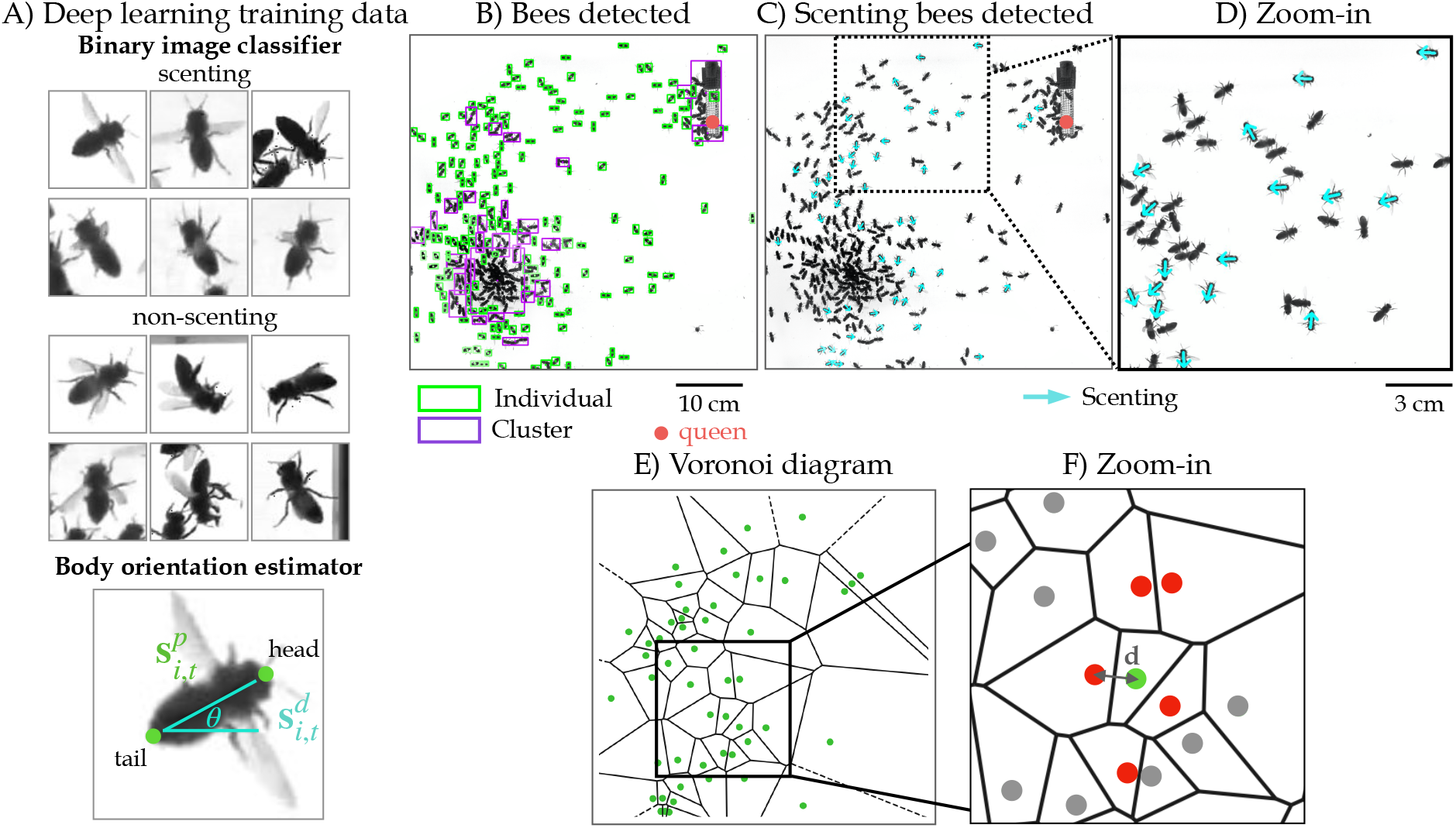
Image Analysis. A) Examples from labeled/annotated data–sets, used to train a CNN models to detect bees and classify them to scenting vs. non–scenting categories as well as estimate their body orientations and therefore scenting directions. B) Example of a snapshot where individual bee are detected. Individual bees are highlighted in green and clusters in purple. No purple box is drawn for large clusters to reduce clutter. C) The same snapshot showing the recognition of the scenting bees (highlighted in cyan). D) A zoomed–in fraction of the snapshot that shows the detected body orientation of the scenting bees, shown by an arrow. E) Example Voronoi diagram of the frame image from B), showing the centroids of the scenting bees as green dots inside cells (black lines). F) A zoomed-in fraction of the diagram in E) that shows a particular scenting bee (green dot) in a cell and neighbor scenting bees (red dots) that occupy the cells adjacent to the green bee. Grey dots represent scenting bees that are not counted as neighbors of the green bee. The Euclidian distance between the bee in green and all the bees in red are computed and averaged for this particular scenting bee.

We trained a convolutional neural network (CNN) to classify individual bee images as scenting or non–scenting, examples shown in Fig. S2A. Neural networks require large annotated datasets. To address this issue, we developed an annotation tool to rapidly generate 28,458 labeled images. The labeled dataset was split into training, validation, and test sets in order to assess the generalizability of the model. Although scenting is a time–dependent behavior, given the characteristic behavior of vigorous wing–flapping in scenting bees, we are able use the wing angle as a proxy for scenting, enabling the use of static images to identify scenting bees. We trained a ResNet–18 model [He et al., 2016] with data augmentation (horizontal and vertical flipping, brightness adjustments, scaling, translation, and rotation) and balanced sampling to combat the class imbalance (9:1 non–scenting to scenting) for 1203 epochs with early stopping to prevent over–fitting. We achieved 95.17% accuracy, 95.39% true positive rate, and 95.15% F1 score (95.39% recall, 94.92% precision) on the test set, indicating that our model can generalize to unseen data during training.

We also employ deep learning for the task of orientation prediction by posing it as a regression problem, allowing the estimation of continuous orientation angles. The same CNN architecture from the classification task is used, with a modified loss function to output continuous values for the predicted angles:

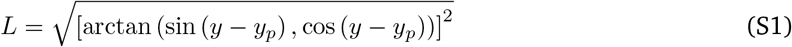

where *y* is the true label, and *y_p_* is the network’s prediction. Similar to the scenting bee classification task, we created a labeled dataset of 15,435 images each with head and tail positions, from which we compute the ground–truth orientation angle (Fig. S2A). On the test set, this model achieves 96.71% with 15° of error tolerance. Together, these computer vision and deep learning methods allow us to automatically process long recordings to detect individual bees (Fig. S2B), classify scenting bees and non–scenting bees, and estimate their body orientations for scenting directions (Fig. S2C–D).

### 5.3 Analysis of extracted data

With the positions and orientations of scenting bees extracted from the recordings, we obtain several measurable time-series properties from the experiments. The number of scenting bees are extracted per frame of each density experiment, presented as a rolling mean with the window size of 100 frames (Fig. 2B). To compute the distance between neighbor scenting bees, for each frame of an experiment, we define a Voronoi diagram whose points are the positions of the scenting bees in that frame. For a given scenting bee, its neighbors are the points that create cells adjacent to the cell of that bee. The Euclidian distance from the bee to its neighbors are computed and averaged, and for a given frame, these distances for all scenting bees are averaged. The time–series data for this property is presented as a rolling mean with a window size of 100 frames (Fig. 2C). The final property we extracted from the experimental data is the average distance to the queen (Fig. 2D). Because the computer vision method to detect individual bees cannot find every single individual bee (when bees touch or overlap, they count as a clustered group), we alternatively computed the average distance of all black pixels to the queen’s location in binary frame images. Because the queen’s cage is stationary and the remaining black pixels in the arena only represent the moving worker bees, this is an effective method to measure this distance to the queen over time.

We note that because the bees were placed in the fridge before the experiments to reduce flying capability for easier handling, they were not yet very active in the initial clusters. Hence, our analysis of the experiments can begin before all bees are spread out to capture the sharp increase in the number of scenting bees at the beginning. Besides the initial peak in the number of scenting bees at the beginning in each of the experiments, two experiments also show a sharp peak in the later half of the recording (Fig. 2B). These later peaks are a result of a sudden mass scenting event with many bee participants, and require further investigation.

### 5.4 Correlation definition

Now that we have the positions and directions of all scenting bees in our experiments, we turn to ask how these scenting events are correlated with the spatiotemporal density of bees in the arena, defined as *ρ*(*x, y, t*). For each scenting bee *i* at time *t*, we define its position as 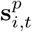, and its direction of scenting as 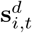 (unit vector). If the scenting bees provide directional information to non–scenting bees, we could treat 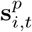 and 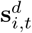 as a set of gradients that define a minimal surface of height *f* (*x, y, t*). Hence, *f* (*x, y, t*) corresponds to the probability that a randomly moving non–scenting bee will end up at position (*x, y*) by following the local scenting directions of scenting bees:

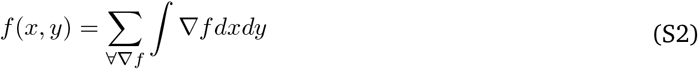

where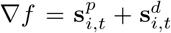. Specifically, we regularize the least squares solution of the reconstruction of a surface from its gradient field, using Tikhonov regularization [Harker and O’Leary, 2008, Harker and O’Leary, 2011]. An example of the reconstruction of the surface is shown in (Fig. 1B). Finally, we define the correlation between the attractive surface *f* (*x, y, t*) and the concentration of bees *ρ*(*x, y, t*′), using mutual information:

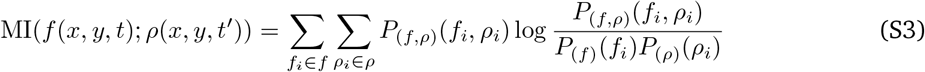

where *P*_(*f, ρ*)_ is the the joint probability mass function of *f* (*x, y, t*) and *ρ*(*x, y, t*′), and *P*_(*f*)_(*f_i_*) and *P*(_*ρ*_)(*ρ_i_*) are the marginal probability mass functions of *f* (*x, y, t*) and *ρ*(*x, y, t*′), respectively.

To scale the results between 0 (no mutual information) and 1 (perfect correlation) for interpretability, we use the normalized mutual information (NMI):

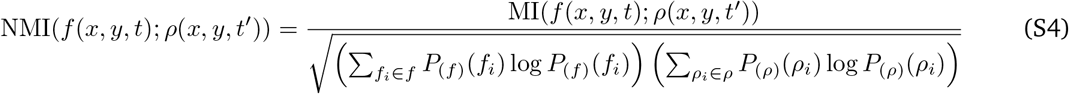

**Figure S3:**
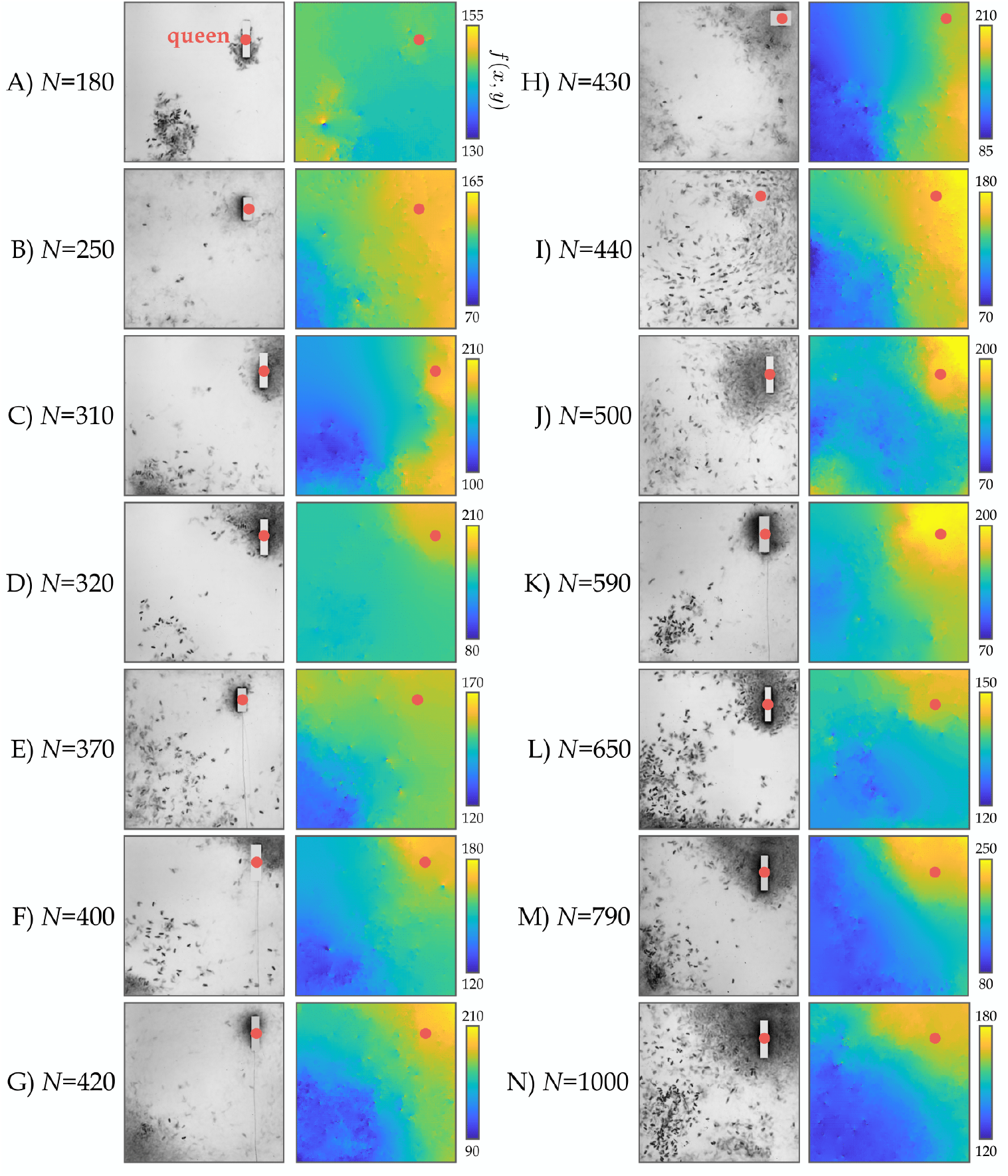
Concentration of Bees vs Attractive Surfaces. A–N) For each of the density experiments, we show the average frame of the last 5 minutes of the experiment when we typically have bees’ final aggregation around the queen, with the corresponding attractive surface averaged over the entire experiment. Note that in the frames, the queen’s cage is replaced by a rectangle that matches the color of the background to not interfere with the mutual information computation. The red dot denotes the queen’s location. The aggregation around the queen typically correlates with the areas of maximal *f* (*x,y*) in the attractive surface. The mutual information is computed for each of these pairs of average frame and average surface.

Our controls for the NMI calculations include generating 10 pairs of uniformly random 1000 × 1000 matrices that perfectly correlate (i.e., they are identical pairs of matrices). Averaged over the 10 trials, the NMI is 1.00 as expected. For the case of highly uncorrelated matrices, we generated 10 pairs of uniformly random 1000 × 1000 matrices, which have a low average NMI of 0.006.

**Table S1:**
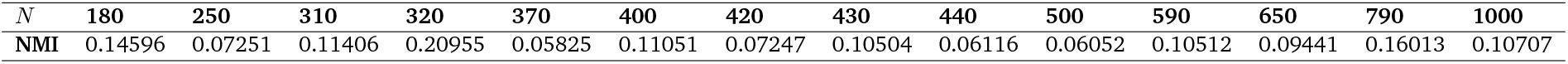
Mutual Information Between Concentration of Bees and Average Attractive Surface. For all density experiments, the mutual information and normalized mutual information between the attractive surface averaged over the entire experiment and the average frame from the last 5 minutes of the experiment are shown.

### 5.5 A candidate set of behavioral rules

To explore the honeybee communication network in the queen–finding context, we built an agent–based–model predicated on a pheromone diffusion and concentration gradient model and a set of simple behavioral rules. We chose this family of models to study how simple behavioral rules can generate complex collective behaviors.

In the agent–based model, we model pheromone diffusion using the 2–dimensional (2–D) diffusion partial differential equation to describe the concentration of pheromone, *C*(*x, y, t*), at a position and time:

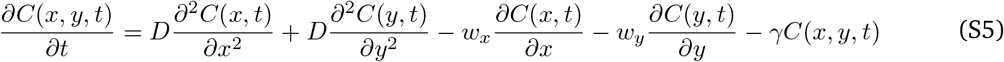

where *C*(*x, y, t*) is the pheromone concentration at position [*x, y*] at time *t, w_x_*, and *w_y_* are the *x* and *y* components of emission vector, respectively, *D* is the diffusion coefficient, and γ is the decay constant. Treating a single scenting bee as a point source of localized and instantaneous pheromone emission, we solve eq. S5:

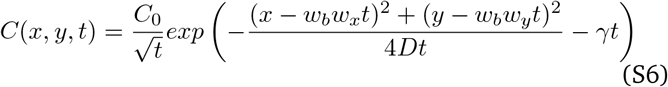

where *C*_0_ is the initial pheromone concentration and *w_b_* is the magnitude of directed emission. The environmental parameters of the model include: the size of the 2–D square arena (*X_min_* and *X_max_*) and the size of a grid cell (*δ_X_*), the start and final time of the simulation (*t_i_* and *t_f_*) and the time integration constant (*δ_t_*).

The queen bee acts as a stationary and frequent source of pheromone and a global point of convergence. Her pheromone emission is modeled as an axi–symmetric diffusion process, i.e., without directional bias (*w_b_* = 0). Worker bees in the simulation obey the following behavioral rules: (1) Workers begin by performing a random walk in the arena. Workers near the queen will detect her pheromone if it is above their detection threshold (T) at their positions in the arena. (2) If the threshold is met, a worker bee will determine her heading direction as the direction uphill the gradient.The negative vector of the gradient scaled by *w_b_* is the direction to emit pheromone and disperse it via wing fanning for signal propagation. The bee will then either walk up the concentration gradient or stand still for a certain time to emit and fan her own pheromones, to inform other bees in the swarm of the queen’s direction, each event with a 0.5 probability. (3) Bees that detect this cascade of secondary signals will follow the same rules to head in the direction of maximum pheromone concentration or emit their own and further propagate the information.

**Figure S4:**
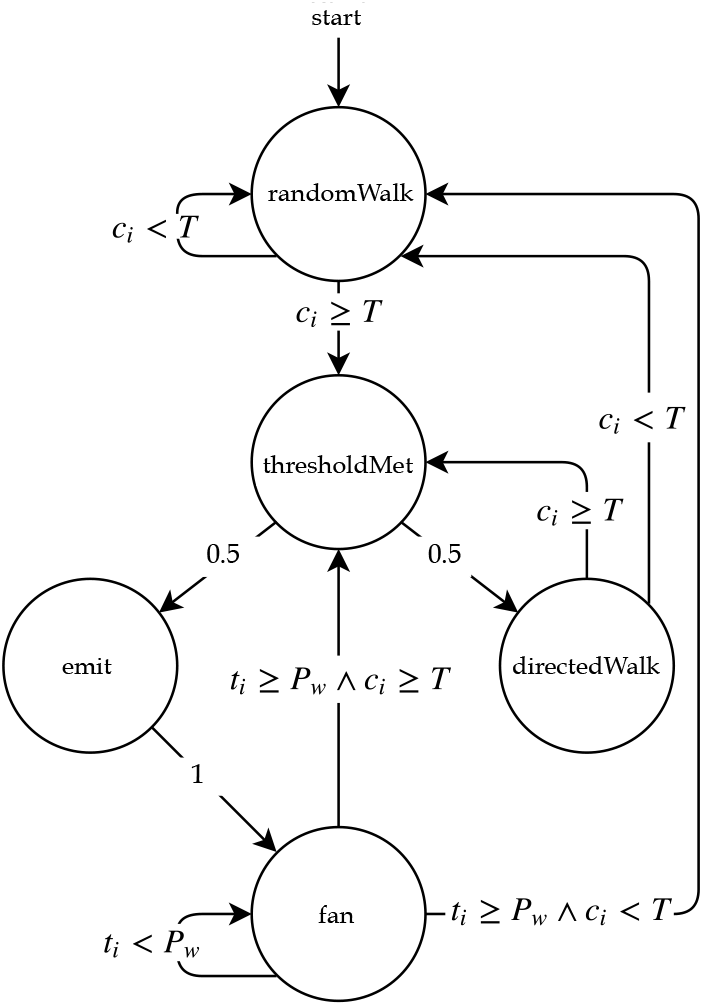
Probabilistic State Machine Diagram. The behaviors of worker bees are presented as a state model with finite states (circles) and probabilistic and conditional transitions (links).

We formalize the worker bees’ behavior as a probabilistic state machine (PSM) [Rabin, 1963]. The PSM consists of a set of finite states that describe bee behavior, such as random walking or fanning, and a probabilistic transition matrix that prescribes the manner in which a bee may change from one state to another. Specifically, the state model *SM_worker_* = (*S, s*_0_,*I, M*), associated with each worker bee in our ABM, defines her set of behavioral rules within the environment, and *SM_worker_* components are fixed across all worker bees and are specified as follows.

- *S* = {*randomWalk, directedWalk, thresholdMet, emit, fan*}, is a set of finite states, where the variable *randomWalk* is a random walk when the activation threshold is not met, *directedWalk* is the walk uphill the concentration gradient, *thresholdMet* is when the concentration threshold of the bee is met, *emit* is the instantaneous release of pheromone, and *fan* is the wing fanning at a constant position.
- *s*_0_ = *randomWalk* is the initial state of each bee
- *I* = {*t_i_, c_i_*}, is a set of flags for the input conditions on state transitions, where for a given bee, *t_i_* is a counter for the time that bee is in the *fan* state and *c_i_* is the concentration at that bee’s position.

For the transition matrix *M*, there are two relevant ABM parameters, *P_w_* and *T*, which represent the emission period comprised of the *emit* and the *fan* state and the concentration threshold over which a worker bee can be activated from state *randomWalk*. The table provides the conditions and probabilities for transitioning from the current state, *s_c_*, to the next state, *s_n_*. The table (Table S2) is defined as follows, with *randomWalk, thresholdMet*, and *directedWalk* abbreviated as *rWalk, tMet*, and *dWalk*, respectively. This state model is also visually represented in a state diagram in Fig. S4.

**Table S2:** Probabilistic State Machine Transition Matrix for Honeybee Behavioral Rules.

| $s_c \backslash s_n$ | rWalk | tMet | emit | fan | dWalk |
| --- | --- | --- | --- | --- | --- |
| rWalk | $c_i < T$ | $c_i \geq T$ | 0 | 0 | 0 |
| dWalk | $c_i < T$ | $c_i \geq T$ | 0 | 0 | 0 |
| tMet | 0 | 0 | 0.5 | 0 | 0.5 |
| emit | 0 | 0 | 0 | 1 | 0 |
| fan | $t_i \geq P_w \wedge c_i < T$ | $t_i \geq P_w \wedge c_i \geq T$ | 0 | $t_i < P_w$ | 0 |

To compute the direction of greatest local change in concentration for a given bee, we calculate the gradient of pheromone concentration:

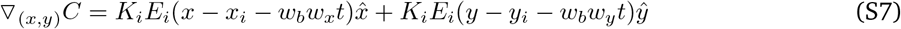

where 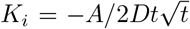 and *E_i_ = exp* (−(*x − x_i_ − w_b_w_x_t*)^2^ + (*y − y_i_ − w_b_w_y_t*)^2^/4*Dt* − γ*t*), *x, y* are the position of the activated bees, and *x_i_, y_i_* are the position of the scenting bees (i.e., pheromone source) *i*. The cumulative gradient for the concentration at a single bee’s position is the sum of the normalized gradients resulting from each pheromone source or emitting bee *i*:

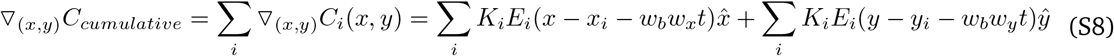

This gradient defines the vector that points in the direction of the bee’s heading for its directed walk. The negative vector of the gradient is the direction for this bee’s pheromone emission for signal propagation, and thus its *x* and *y* components make up the *w_x_* and *w_y_* terms of Eq. S6.

**Table S3:** Agent-based Model Parameters. Free parameters of the ABM are bolded.

| Parameter | Meaning | Value(s) |
| --- | --- | --- |
| $X_{min}, X_{max}$ | Size of grid arena | -3, 3 |
| $\delta_X$ | Grid cell size | 0.01 |
| $t_i, t_f$ | Start and final time | 0, 50 |
| $\delta_t$ | Time integration constant | 0.005 |
| $D$ | Diffusion coefficient | 0.6 |
| $\gamma$ | Decay constant | 108 |
| $(x_q, y_q)$ | Queen 2-D position | (0,0) |
| $P_q$ | Queen emission frequency | 80 |
| $C_{0q}$ | Queen initial concentration | 0.0575 |
| $w_{bq}$ | Queen bias scalar | 0 |
| $(x_w, y_w)$ | Worker 2-D position | $\in [X_{min}, X_{max}]$ |
| $P_w$ | Worker emission period | 80 |
| $C_{0w}$ | Worker initial concentration | 0.0575 |
| $w_{bw}$ | **Worker bias scalar** | 0, 10, 20, 30, 40, 50, 60 |
| $N_w$ | **Worker density** | 50, 100, 300 |
| $T$ | **Worker activation threshold** | 1e-4, 1e-3, 0.01, 0.025, 0.05, 0.075, 0.1, 0.20, 0.25, 0.3, 0.4, 0.5, 0.6, 0.7, 1.0 |

### 5.6 Construction of the phase diagrams

We extracted several straightforward properties from the model data for time-series analyses presented in Fig. 4A–B, D: the bees’ average distance to the queen, the average number of scenting bees, and the average distance to the queen from the farthest active bee. To obtain the clustering data to characterize the growth of the queen’s cluster size and the number of clusters that form over time (Fig. 4C, S5A–B), we use the density-based spatial clustering of applications with noise (DBSCAN) algorithm (eps: 0.25, minimum number of bees to form a cluster: 5) to cluster bees at every time step [Ester et al., 1996]. Finally, to find the time when the average distance to the queen reaches a plateau value (Fig. 4E), we find the time step after which the difference between contiguous distance values is 0.001 or below for at least 300 time steps.

**Figure S5:**
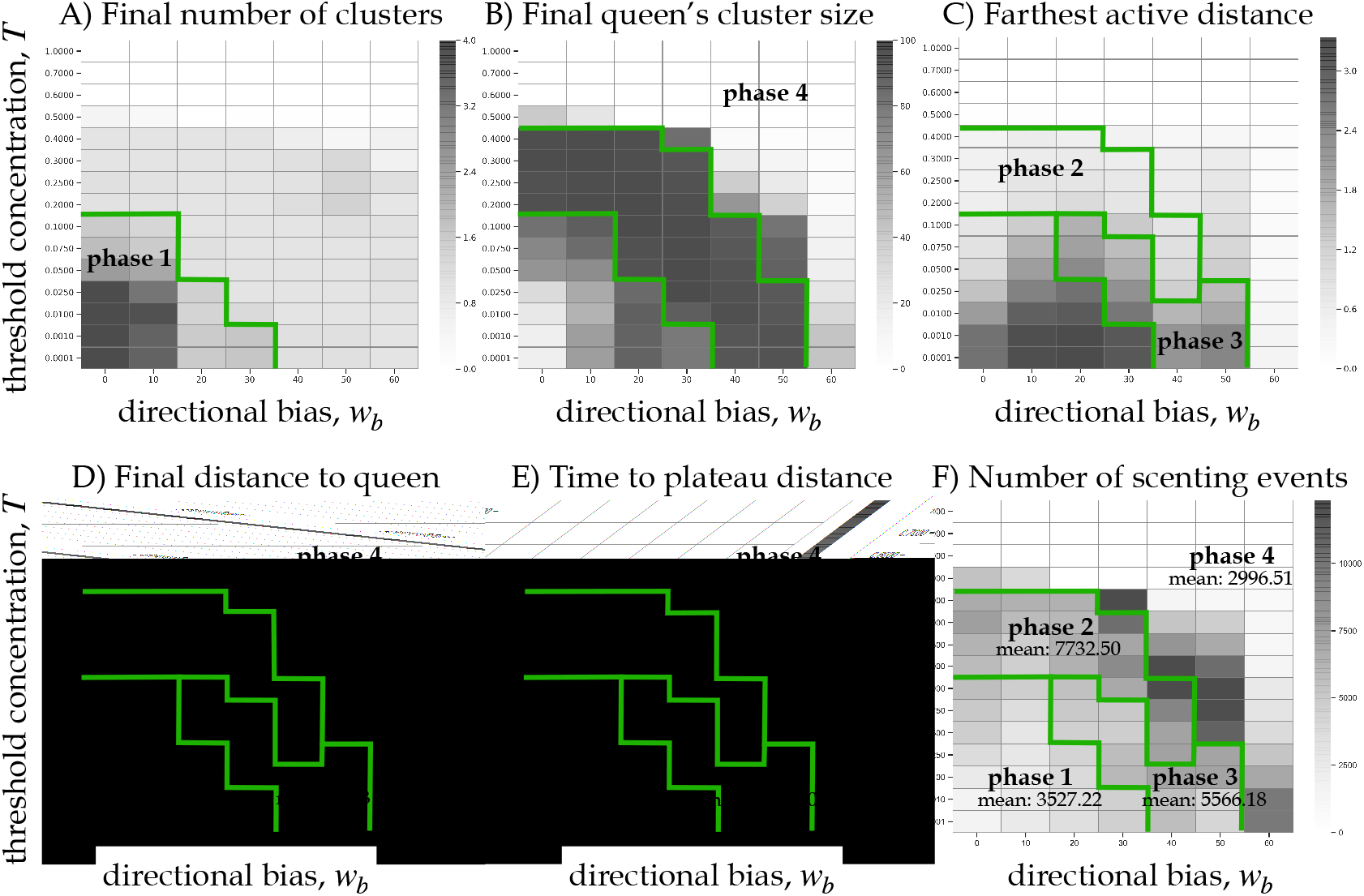
Summary Heatmaps for N = 100. A–C) Summary heatmaps used to construct the phase diagrams. A) The average final number of clusters as a function of *T* and *w_b_*, used to identify phase 1 simulations that result in more than 1.4 clusters. B) The average final size of the queen’s cluster, used in combination with A) to identify phase 4. Phase 4 simulations have 0–1.4 clusters and the cluster size is less than 90 bees. C) The average distance of the farthest active bee to the queen during the initial 1000 time steps of the simulation, used to distinguish between the random walk in phase 2 and percolation in phase 3. Phase 2 simulations must have this average distance below 1.5, while simulations in phase 3 must meet the threshold of 1.5 or greater. D–F) Summary heatmaps showing various properties of each phase, with average values for each phase annotated and all phases overlaid. D) The average final distance to the queen. E) The average time the simulations reach a plateau distance to the queen. F) The average total number of scenting events up to the time to the plateau distance.

We then sum the number of scenting events from the beginning of the simulation to the time to the plateau distance to the queen (Fig. 4F).

To construct the phase diagrams, we use three summary heatmaps to distinguish the phases: the final number of clusters, the final queen’s cluster size, and the distance of the farthest active bee to the queen (Fig. S5A–C). For trials with the intermediate density of bees, (*L*^2^/100), we sequentially applied the following conditions to each simulation to group it into the appropriate phase:

- If the final number of clusters > 1.4: Phase 1
- If the final number of clusters 0 – 1.4 and the final queen’s cluster size < 90 bees: Phase 4
- If the farthest active distance < 1.5: Phase 2
- If the farthest active distance ≥ 1.5: Phase 3

For low density of bees, (*L*^2^/50), the sequential conditions are:

- If the final number of clusters > 1.1: Phase 1
- If the final number of clusters 0 – 1.1 and the final queen’s cluster size < 45 bees: Phase 4
- If the farthest active distance < 1.2: Phase 2
- If the farthest active distance ≥ 1.2: Phase 3

Lastly, for high density of bees, (*L*^2^/300), the sequential conditions are:

- If the final number of clusters > 2.0: Phase 1
- If the final number of clusters 0 – 2.0 and the final queen’s cluster size < 270 bees: Phase 4
- If the farthest active distance < 2.3: Phase 2
- If the farthest active distance ≥ 2.3: Phase 3

**Algorithm 1.**
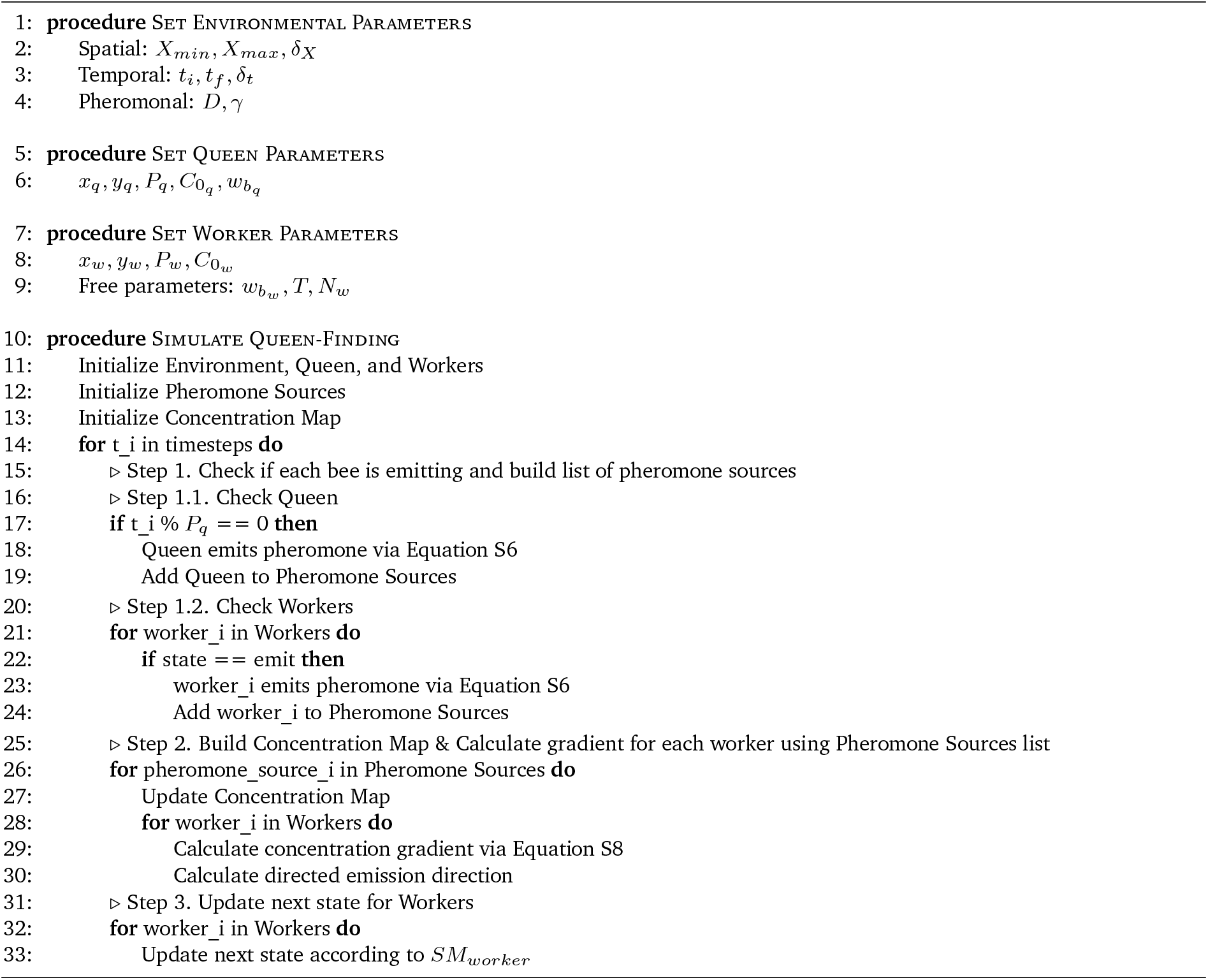
Honeybee Queen-Finding Algorithm

## Supplementary Movies

- S1: Example closeup movie of a scenting bee (left panel) and a non-scenting bee (right panel) in the semi-2D experimental arena from an aerial view. The scenting bee is pointing up her abdomen and fanning vigorously. The non-scenting bee is standing still.
- S2: Example movie of results from the image analysis pipeline as shown in Fig. S2. The left panel shows the detection of individuals (green boxes) and clusters (purple boxes) using the computer vision methods of adaptive thresholding, morphological transformations, and connected components. Note that for very large clusters, no purple boxes are drawn to reduce visual clutter. The right panel shows the recognition of scenting bees and the estimation of their scenting directions (cyan arrows) using convolutional neural networks.
- S3: Example movie of the *N* = 320 experiment and data shown in Fig. 1. The left panel shows the experimental movie with annotated scenting directions on scenting bees (cyan arrows). The center panel shows the attractive surface reconstructions *f* over time to show the correlation between scenting directions and aggregation location of the bees. The right panel shows the time-series data of the number of scenting bees over time, where we observe a sharp peak at the beginning and gradual decrease.
- S4, S5, S6: Three example movies of the experiments of various densities (*N* = 320,500,790) shown in Fig. 2. The queen bee is in a cage at the top right corner of the arena. Worker bees are placed at the bottom left corner at the beginning. Over time, they collectively form a communication network using the scenting behavior and end up aggregating around the queen.
- S7: Example movie of a simulation in phase 1 in the agent-based model (*w_b_* = 0, *T* = 0.01, *N* = 300, *C*_0_ = 0.0575, *D* = 0.6, γ = 108). In this phase at relatively low values of T and *w_b_*, the bees aggregate into small clusters that are homogeneously spread throughout the simulation box.
- S8: Example movie of a simulation in phase 2 in the agent-based model (*w_b_* = 10, *T* = 0.5, *N* = 300, *C*_0_ = 0.0575, *D* = 0.6, γ = 108). In this phase at relatively higher values of *T* and *w_b_*, only bees that randomly walk near the queen are activated (i.e., scent or perform the directed walk up the gradient) and join the local cluster around her. This is similar to a Diffusion Limited Aggregation that results in a sparse fractal-like cluster.
- S9: Example movie of a simulation in phase 3 in the agent-based model (*w_b_* = 30, *T* = 0.05, *N* = 300, *C*_0_ = 0.0575, *D* = 0.6, γ = 108). In this optimal phase at relatively low values of *T* and high values of *w_b_*, the activated bees create a percolating network of senders and receivers of the pheromone signals. The aggregation process and the growth of the queen’s cluster is the fastest compared to other phases.
- S10: Example movie of a simulation in phase 4 in the agent-based model (*w_b_* = 0, *T* = 1.00, *N* = 300, *C*_0_ = 0.0575, *D* = 0.6, γ = 108). In this phase, the high activation threshold *T* results in no workers bees ever activated and no clusters formed around the queen. The bees perform the random walk throughout the entire simulation.

## Code

Code for the image analysis of the experiments and the agent-based model will be made available upon publication.

## Author Contributions

O.P. designed and directed the project. D.M.T.N. and A.M. performed the experiments. A.M. performed preliminary analysis of the experimental data. D.M.T.N. and M.L.I. developed the computer vision pipeline for image analysis in discussion with K.B. and G.J.S.. D.M.T.N. and O.P. analyzed the experimental data. O.P., D.M.T.N., and M.L.I. designed and coded the agent-based model. M.L.I. derived Equation S8. D.M.T.N. and O.P. analyzed the model results. M.L.I. helped write the image analysis methods. D.M.T.N. and O.P. wrote the manuscript with feedback from all authors.

## Acknowledgments

This work was supported by the BioFrontiers Institute at the University of Colorado Boulder (internal funds) and by the National Science Foundation Graduate Research Fellowship under Grant No. DGE 1650115 (D.M.T.N.). Any opinion, findings, and conclusions or recommendations expressed in this material are those of the authors(s) and do not necessarily reflect the views of the National Science Foundation. We also acknowledge funding from OIST Graduate University (K.B., G.J.S.) and from Vrije Universiteit Amsterdam (G.J.S.). We thank Seneca Kristjonsdottir and Christopher Borke for bee management; Gary Nave and Michael Neuder for assistance in developing image analysis pipeline; Aubrey Kroger and Emily Walker for annotating images; Raphael Sarfati for reading and commenting on the manuscript. We thank Prof. L. Mahadevan, Prof. Massimo Vergassola, Prof. Gene E. Robinson, Prof. Olav Rueppell, and members of the Peleg lab for insightful feedback and discussions.

